# Deletion of an enhancer in FGF5 is associated with ectopic expression in goat hair follicles and the cashmere growth phenotype

**DOI:** 10.1101/2020.08.24.264754

**Authors:** Yefang Li, Shen Song, Xuexue Liu, Yanli Zhang, Dandan Wang, Xiaohong He, Qianjun Zhao, Yabin Pu, Weijun Guan, Yuehui Ma, Lin Jiang

## Abstract

Research on cashmere growth has a significant effect on the production of cashmere and a profound influence on cashmere goat breeding. Whole-genome sequencing is a powerful platform to rapidly gain novel insights into the identification of genetic mechanisms underlying cashmere fiber growth. Here, we generated whole-genome sequences of 115 domestic goats from China, Nepal and Pakistan, including 51 cashmere goats and 64 non-cashmere goats. We found genetically distinct clusters according to their geographic locations but genetic admixture or introgression may have occurred between the Chinese and Nepalese goats. We identified that the *fibroblast growth factor 5* gene (*FGF5*) shows a strong signature for positive selection in the cashmere goat. The 505-bp indel variant at the *FGF5* gene locus appeared to be strongly associated with cashmere growth. Functional validation showed that the insertion variant may serve as an enhancer for transcription factor binding, resulting in increased transcription of the upstream *FGF5* gene in non-cashmere goats. Our study provides useful information for the sustainable utilization and improved conservation of goat genetic resources and demonstrates that the indel mutation in the *FGF5* gene could potentially serve as a molecular marker of cashmere growth in cashmere goat breeding.

**Author summary:** Cashmere goats have been selected for thousands of years and have become economically significant livestock in China and other central Asian countries. The mechanism of cashmere growth is not well understood because most studies have focused on the investigation of candidate genes. Here, we conducted a comprehensive whole-genome analysis for selection signatures in a total of 115 goats from 15 genetically diverse goat breeds. The results revealed a strong selection signature at the *FGF5* gene locus associated with the cashmere growth phenotype. A 505-bp indel was located in the downstream region of *FGF5* and significantly separated in the cashmere goats versus non-cashmere goats. Functional effect analysis of the indel revealed that it may act as an enhancer by specifically binding transcription factors to mediate quantitative changes in *FGF5* mRNA expression. Our study illustrates how a structural mutation of the *FGF5* gene has contributed to the cashmere growth phenotype in domestic goats.

## Introduction

Cashmere wool, usually simply known as cashmere, is a fiber obtained from cashmere goats, pashmina goats, and some other breeds of goat. This fiber has been used to make yarn, textiles and clothing for hundreds of years. Cashmere is closely associated with the Kashmir shawl; the word “cashmere” is derived from an anglicization of Kashmir, which occurred in the 19th century when the Kashmir shawl reached Europe from Colonial India [1]. Common usage defines the fiber as wool, but it is finer, stronger, lighter, softer and approximately three times more insulating than sheep wool [2].

Cashmere has been manufactured in China, Mongolia, Nepal and Pakistan for thousands of years. China has the largest number and richest variety of cashmere goats, such as the Inner Mongolia cashmere, Liaoning and Tibetan varieties, and has become the largest producer of raw cashmere, estimated at 15,438 metric tons (in hair) per year [3]. Nepal has a sizeable indigenous goat population with many nondescript goats. Nepalese goat breeds exhibit enormous variations in fecundity; meat, milk and fibre production; disease resilience; and nutritional requirements. Pakistan is the fourth largest goat-producing country after China, India and Nigeria (FAOSTAT, http://www.fao.org/faostat). The major purposes of Pakistani goats are milk, meat and hair [4]. Moreover, some studies have suggested that a second domestication event for cashmere breeds took place in Pakistan [5].

Cashmere goats grow a double coat composed of the guard hair produced by primary hair follicles (PHFs) and the cashmere produced by the secondary hair follicles (SHFs) [6,7]. The staple length and diameter of hair fibers are the main indicators used to evaluate the value of cashmere. Therefore, identification of related genes and molecular mechanisms that regulate cashmere traits is of great significance. In recent years, based on the goat reference genome, several studies have attempted to characterize genetic variations of cashmere fiber traits in different goat populations using a whole-genome sequencing strategy. For instance, the genes *PRDM6, FGF5* [8]*, LHX2, FGF9, WNT2* [9]*, SGK3, IGFBP7, OXTR* [10], and so on, showed a strong selection signature for cashmere growth and length in Chinese goat populations. However, to our knowledge, few studies have identified the causative mutations of the goat *FGF5* gene that underlie cashmere growth in goats. Moreover, the sample size in cashmere goat studies has been limited to Chinese goat breeds, which may not comprehensively analyze cashmere traits.

Here, we sequenced the whole genomes of 115 goats representing 15 breeds from habitats in China, Nepal and Pakistan. To identify the genetic basis for cashmere growth trait in cashmere goats, we performed genomic analysis of selection signatures of goats and identified that the *FGF5* gene showed some of the strongest signatures for positive selection in the cashmere goat genome. Further exploration of the *FGF5* genotypes and functional validation assays indicated that a 505-bp indel mutation located downstream from *FGF5* gene may act as an enhancer, resulting in increased mRNA expression of the *FGF5* gene; moreover, deletion of this enhancer is strongly associated with cashmere growth in cashmere goats.

## Results

### Genomic variants

We used the Illumina HiSeq platform to generate whole-genome sequencing data of 115 goats sampled from China, Nepal and Pakistan (**Fig 1A**). The median genome coverage achieved across the full data set was ~5X (min=2.11X, max=7.78X), representing ~87.44% (min=78.54%, max=89.88%) base coverage per individual genome (**S2 Table**). The alignment ratio of reads to the genome was 93.57%-99.93% (**S2 Table**). Strict read alignment and genotype calling procedures allowed us to obtain a total of 17,534,538 single nucleotide polymorphisms (SNPs). The majority of the autosomal SNPs were located within intergenic (11,821,678, 45.847%) and intronic (11,347,329, 44.007%) regions, with only 0.858% (221,209) located in exonic regions. Approximately 9% were present in downstream or upstream gene regulatory regions. A total of 61,164 missense SNPs and 126,811 synonymous SNPs resulted in a nonsynonymous/synonymous ratio of 0.482 (**S3 Table**).

**Fig 1.**
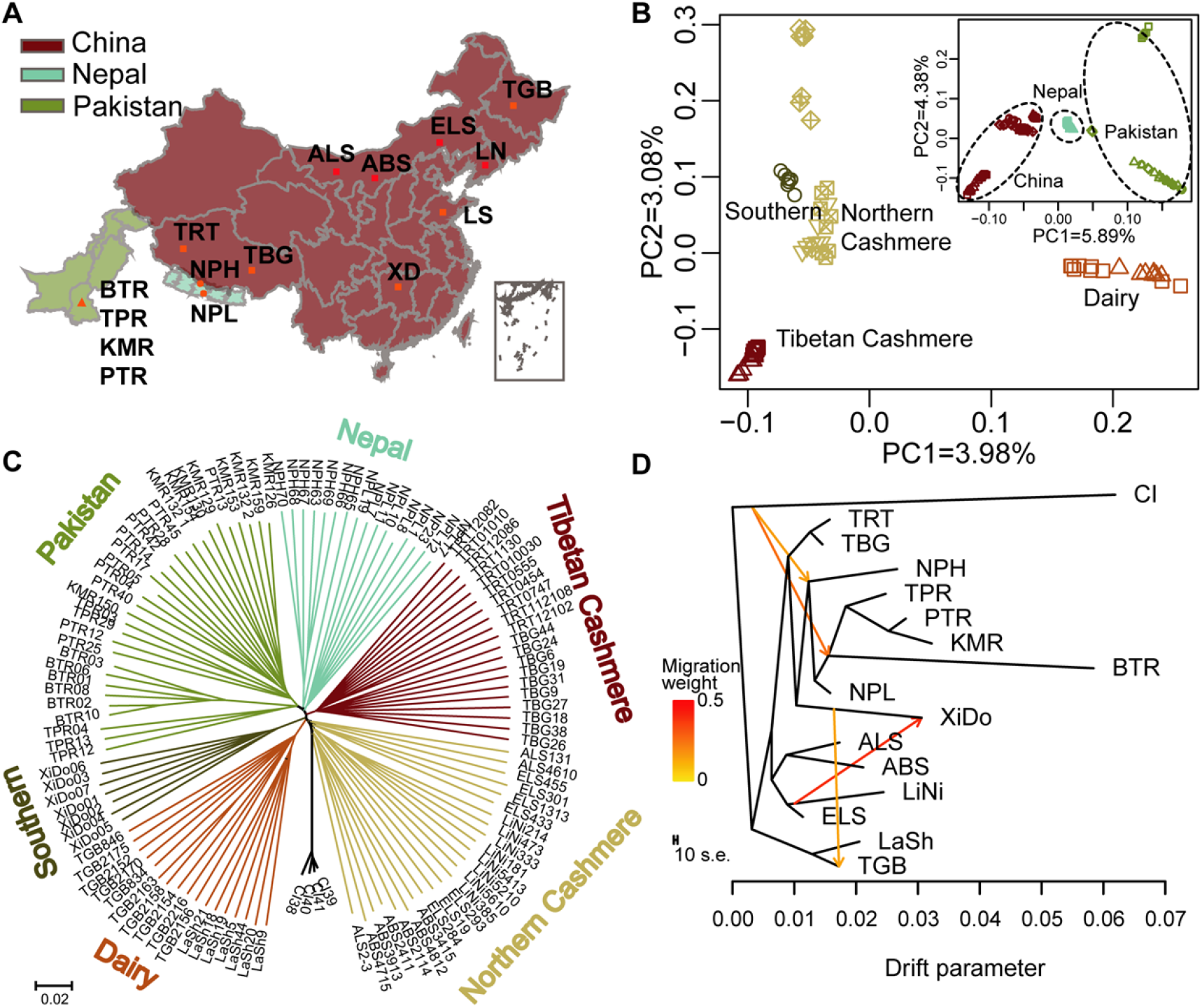
Geographic distribution, genetic structure of Chinese, Nepalese and Pakistani goat breeds. (A) The geographic distribution of 15 goat populations. The red color represent Chinese goats (TGB, Toggenburg dairy goat from Heilongjiang; LaSh, Laoshan dairy goat from Shandong; LiNi, Liaoning cashmere goat from Liaoning; ABS, Arbus cashmere goat; ELS, Erlangshan cashmere goat; ALS, Alashan cashmere goat from Inner Mongolia; TBG, Tibetan Bangor cashmere goat; TRT, Tibetan Ritu cashmere goat from Tibet; XiDo, Xiangdong black goat from Hunan). The blue color represents Nepalese goats (NPH, Nepalese Highland goat; NPL, Nepalese Lowand goat). The green color represents Pakistani goats (BTR, Bugi Toori goat; KMR, Kamori goat; PTR, Pateri goat; TPR, Tapri goat). (B) PCA plots of the first two components of all goats (inner plot) and Chinese goats (outer plot). The fraction of the total variance explained is reported on each individual axis between parentheses. (C) The neighbor-joining tree of the goat breeds, with *Capra ibex* as the outgroup. Bootstrap reported was close to 100%. (D) The ML-TreeMix tree of all goats, with *Capra ibex* as the outgroup, assuming four migration events. Migration arrows are colored according to their weights. Horizontal branch lengths are proportional to the amount of genetic drift parameter that has occurred on the branch. The drift parameter measures the variance in allele frequency estimated along each branch of the tree. The yellow and orange lines indicate the instantaneous admixtures, whereas arrows denote continuous (unidirectional) genes flow.

### Genetic diversity

Compared to other goats, Tibetan cashmere goats were found to show the highest genome-wide heterozygosity levels, fewer runs of homozygosity (ROHs) and the lowest linkage disequilibrium (LD) decay. However, among northern Chinese cashmere goat breeds, Liaoning cashmere goat (LiNi), Alashan cashmere goat (ALS) and Arbus cashmere goat (ABS), exhibited lower genome-wide heterozygosity, more ROHs and more LD levels, while Erlangshan cashmere goat (ELS) exhibited the opposite. This result may be related to more intensive selection breeding. Imported dairy goats and Xiangdong black (XiDo) goat showed similarly high levels of genetic diversity. The genome-wide heterozygosity levels, ROHs and LD decay were found to be lower in Nepalese highland goats but higher in Nepalese lowland goats. Compared to Chinese and Nepalese goat breeds, the Pakistani goat breeds showed less level of genetic diversity, especially the Bugi Toori goat, which is likely a consequence of its inbreeding history [11] (**S4 Table, S1 and S2 Figs**).

### Phylogenetic analyses

Principal component analysis (PCA) of ~3.4 million unlinked SNPs revealed that the Chinese, Nepalese and Pakistani goats can be separately clustered by the first PCA axis. One Pakistani goat breed (BTR) was genetically more distant from other goat breeds (**Fig 1B, upper panel**). Restricting the analysis to Chinese goat breeds revealed four major clusters (**Fig 1B, main panel**), with the first PCA axis separating the Toggenburg (TGB) and Laoshan (LaSh) dairy goat populations, the second PCA axis separating the Tibetan cashmere goat breeds, and the third PCA axis separating the XiDo black goat from southern China (**S3 Fig**). PC1, PC2 and PC3 were able to explain genetic differences of 3.98%, 3.08% and 2.90%, respectively. The admixture analysis results were largely consistent with the PCA results as well as the similar genetic makeup among the Chinese, Nepalese and Pakistani goats (**S4 Fig**). When K=4, the dairy goats, Chinese goats, BTR goat and the remaining goats were genetically distinct; when K=6 and K=7, the Chinese native goats were divided into the XiDo goat from southern Chinese, the northern Chinese cashmere goats and the Tibetan cashmere goats.

We next investigated the ML-TreeMix tree [12] and the distance-based neighbor-joining tree [12] using *Capra ibex* as the outgroup among all the goats. The distance-based neighbor-joining tree displayed six clades according to location or specific goat trait, which were consistent with the PCA and admixture analysis results (**Fig 1C**). The reliability of the neighbor-joining tree was estimated by 100 bootstrap pseudoreplicates. The ML-TreeMix tree without migration events (ML=0) inferred from the TreeMix analysis divided the 115 goats into six clusters, which were consistent with the neighbor-joining tree results (**S5A Fig**). When M=1 and M=2, the results suggested that gene exchange occurred between wild and Nepalese and Pakistani goats (**S5B and S5C Fig**); when M=3, genetic materials of LiNi flowed to XiDo (S5D Fig); and when M=4-6, genes flowed from XiDo to the TGB, LaSh and Nepalese lowland (NPL) goat breeds (**Fig 1D, S5D-S5F Fig**).

### Genome-wide selection scans for cashmere growth

To detect the positive selection signatures within 100 kb sliding windows, we next scanned the cashmere goat (including TBG, TRT, LiNi, ABS, ELS, ALS and NPH) genomes relative to non-cashmere goat (including TGB, LaSh, XiDo, NPL, BTR, KMR, PTR and TPR) genomes by using three statistical methods, namely, *F*_ST_, θ_π_ ratio (θ_π-noncash_/θ_π-cash_) and ZHp (**Fig 2**). The top-5% selection candidates that were common to all three statistical methods identified 982 windows, which annotated a total of 378 protein-coding genes (**S5-S8 Tables, S6 Fig**). Enrichment analyses for Gene Ontology terms revealed that the melanocortin receptor activity (GO:0004977, *P*-value = 2.33E-06), response to stimulus (GO:0050896, *P*-value = 4.70E-07), developmental process (GO:0032502, *P*-value = 2.07E-06) and cellular metabolic process (GO:0044237, *P*-value = 1.07E-06) categories were significantly overrepresented (**S9 Table**). Notably, the genome window containing the *FGF5* locus was under higher selection (chromosome 6, **Fig 2**). This gene is known to be involved in hair growth. Moreover, several genes under strong selection signals are plausibly related to metabolism, inflammation, melanin precipitation and high-altitude adaptation. For example, *STIM1* is an endoplasmic reticulum calcium sensor involved in regulating Ca2+ and metabotropic glutamate receptor signaling in the nervous system [14,15]. The CERT protein mediates the START pathway of ceramide transport in a nonvesicular manner and the amphiphilic cavity of the START domain is optimized for specific binding of natural ceramides [16,17]. *NOP14* plays significant roles in the proliferation and migration of pancreatic cancer cells [18]. The mutations in the *SGCB* gene can lead to a loss of functional protein and result in limb-girdle muscular dystrophy disease. MYCBP2 is a member of the PHR protein family and an E3 ubiquitin ligase, and it was shown to have important functions in developmental processes, such as axon termination and synapse formation [20]. *WARS2* has low enzyme activity, and inhibition of *WARS2* in endothelial cells reduces angiogenesis [21,22]. *MC1R* and *KIT* genes have been implicated in human and animal hair pigmentation, reflecting a role in the development and function of melanocytes [23,24]. The *DSG3* gene is responsible for the high-altitude adaptation of the Tibetan goat [25,26].

**Fig 2.**
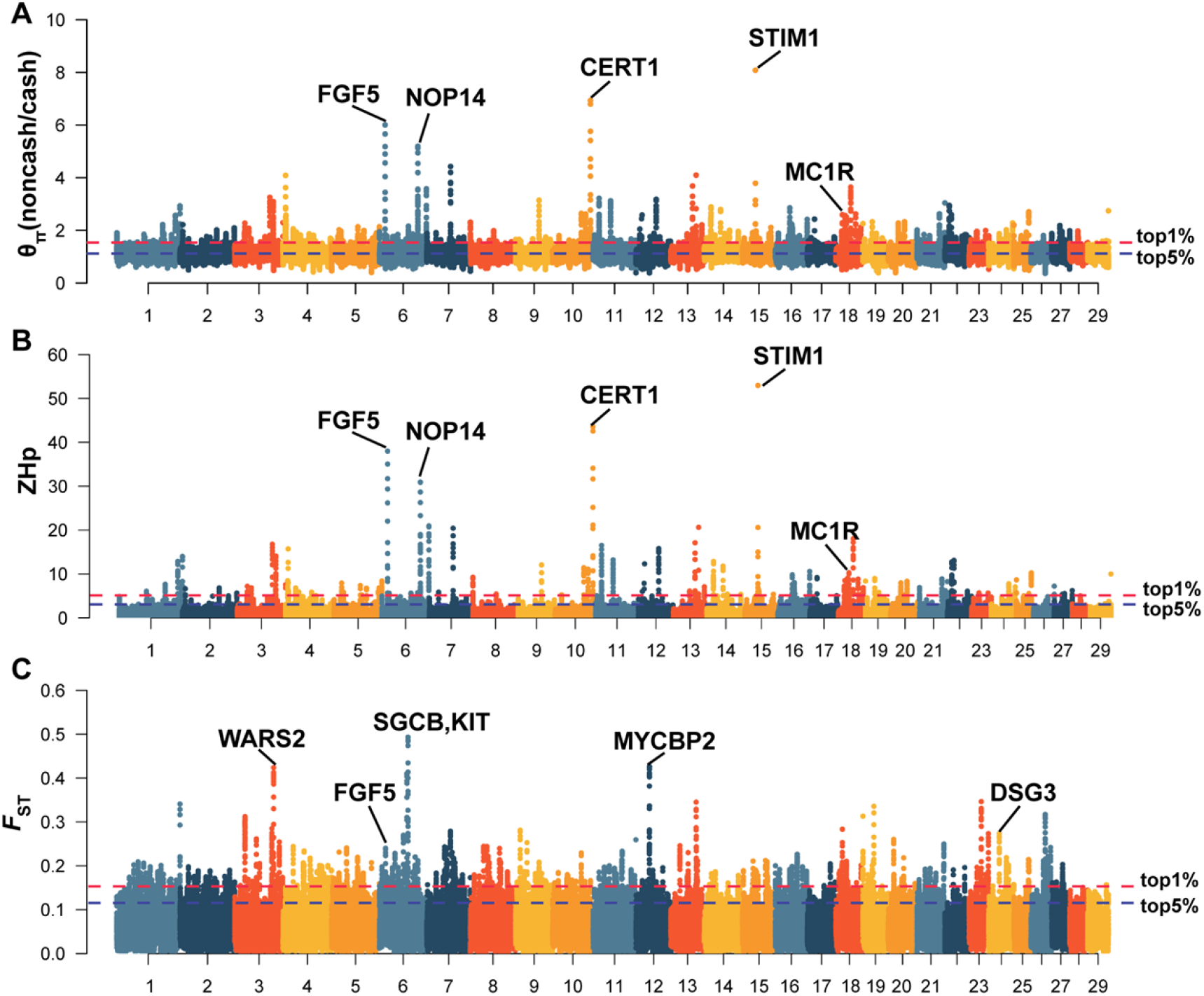
Positive selection scans for cashmere growth. Cashmere goats are compared with non-cashmere goats. The nucleotide diversity θπ ratio (θπ-noncash/θπ-cash) (A), the transformed heterozygosity score ZHp (B) and the population genetic differentiation *F*_ST_ values (C) are calculated within 100 kb sliding windows (step size 10 kb). The significance threshold of a selection signature was arbitrarily set to the top 5% percentile outliers for each individual test and is indicated with blue horizontal dashed lines. The red horizontal dashed lines delineate the top 1% quantile.

### Annotation of variants under positive selection in *FGF5*

We next sought to further refine the selection targets within the *FGF5* locus by using three different methods, namely, the θ_π_ ratio (θ_π-noncash_/θ_π-cash_), Tajima’s D and *F*_ST_, and detecting the read depth. We noticed a 505-bp deletion within the most significant selection region in the *FGF5* gene (**Fig 3, S10-S13 Tables**), located at position 95,454,689-95,455,189 of chromosome 6 (**Fig 4A**). The deletion variant is present in cashmere goats, suggesting that it arose in an independent genetic background.

**Fig 3.**
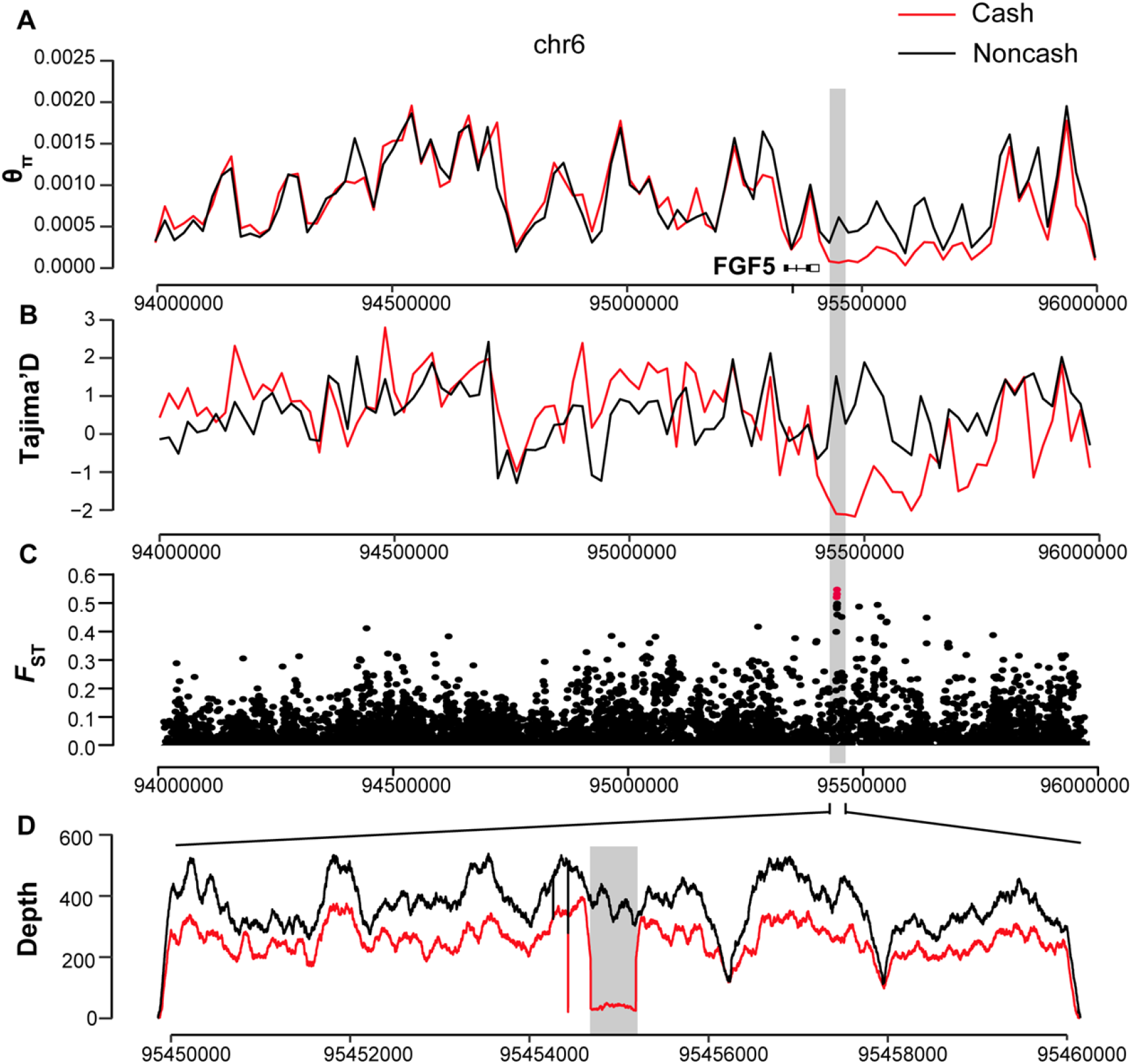
The strongest positive selection signature around the *FGF5* peak. The θ_π_ ratio (A), Tajima’s *D* (B) and *F*_ST_ value (C) are plotted against the peak position from 94.0 Mb to 96.0 Mb, and the read depth value (D) is plotted against the peak position from 95.45 Mb to 95.46 Mb on chromosome 6. Both θ_π_ ratio and Tajima’s *D* values were based on a 20 kb window and a 20 kb step. The red and the black lines represent cashmere and non-cashmere goats, respectively. The gray columns represent the strongest positive selection signature in the region considered. The small black boxes and short lines represent the gene structure of *FGF5*.

**Fig 4.**
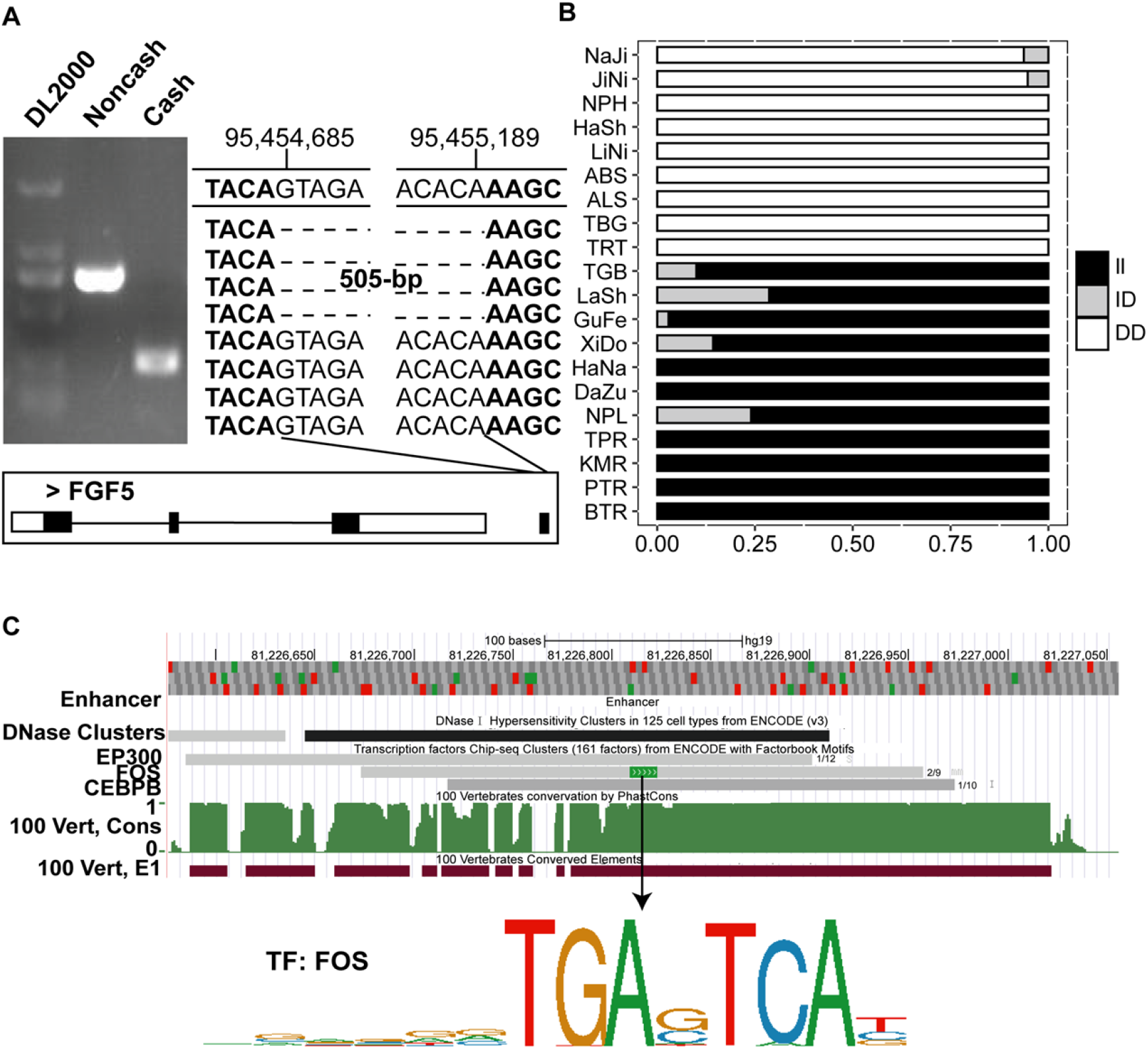
Annotation of the indel variant in the *FGF5* gene showing positive selection signatures. (A) The PCR amplification of 505-bp indel variant, generating a 267-bp fragment in all cashmere goats while a 772-bp fragment in non-cashmere goats. The indel is located at position 95,454,689-95,455,189 of chromosome 6 in the downstream of *FGF5* gene. (B) Genotypes of indel were determined in a larger population (*N* = 288 goats). II represents homozygous insertion genotype; ID represents heterozygous indel genotype; DD represents homozygous deletion genotype. Cashmere goat breeds include NaJi (Nanjiang cashmere goat), JiNi (Jining grey goat), NPH (Nepalese highland goat), HaSh (Hanshan white cashmere goat), LiNi (Liaoning cashmere goat), ABS (Arbus cashmere goat), ALS (Alashan cashmere goat), TBG (Tibetan Bangor cashmere goat) and TRT (Tibetan Ritu cashmere goat). Non-cashmere goat breeds include TGB (Toggenburg dairy goat), LaSh (Laoshan dairy goat), GuFe (Guangfeng goat), XiDo (Xiangdong black goat), HaNa (Hainan black goat), DaZu (Dazu black goat), NPL (Nepalese lowland goat), BTR (Bugi Toori goat), KMR (Kamori goat), PTR (Pateri goat) and TPR (Tapri goat). (C) The insertion fragment of *FGF5* gene in humans contains a highly conserved FOS transcription factor binding site (TGAGTCA) in the UCSC database.

We designed primers spanning the breakpoint of the deletion to genotype the indel variant by gel electrophoresis, generating a 267-bp fragment in all cashmere goats but a 772-bp fragment in non-cashmere goats (**Fig 4A**). To further identify the possible functional consequences of the deletion variant, we investigated whether it showed any association with cashmere growth, extending our analysis to a more comprehensive panel of 288 goats originating from 20 populations (**S14 Table**). The results confirmed a remarkable correlation between the frequencies of the indel variant and cashmere growth. Cashmere goat breeds showed the highest allelic frequencies of deletion (>0.9), whereas non-cashmere goat breeds showed higher allelic frequencies of insertion (nearly 0.8, **Fig 4B**).

Furthermore, the insertion fragment of the *FGF5* locus in humans showed a high conservation score in the 100-vertebrate animals alignment (e.g., goat, mouse, cat, dog, sheep, yak and donkey); by using UCSC database, we also found that the insertion fragment contains EP300, FOS and CEBPB transcription factors among different species [27,28] (**Fig 4C**). This finding indicated that the indel fragment has cis-regulatory effects for *FGF5* gene transcription.

### Biological significance of the indel variant

Since the indel variant contains an extremely conserved FOS transcription factor binding site, we sought to functionally verify the effect of the binding site in this variant. A pair of biotin-labeled probes were designed, namely, a wildtype probe containing TGAGTCA and a mutant probe excluding the site. After the binding reaction with nucleoprotein fractions from NIH/3T3 cells derived from mouse embryonic fibroblasts, the wildtype probe resulted in an obvious positive protein complex band but the mutant probe did not. The addition of either 80-fold or 160-fold cold probes to the reaction system produced a significantly thinner binding band and significantly weakened grayscale compared to those with wildtype probe. The weaker binding reaction may be due to the lower cold probe concentration or higher nucleoprotein concentration making the competitive reaction incomplete. After adding c-FOS antibody to the reaction system, a complex was obviously retained at the top of the gel (**Fig 5A**). We thus hypothesized that the indel variant can specifically bind to the FOS transcription factor, which may have important regulatory effects on upstream target genes.

**Fig 5.**
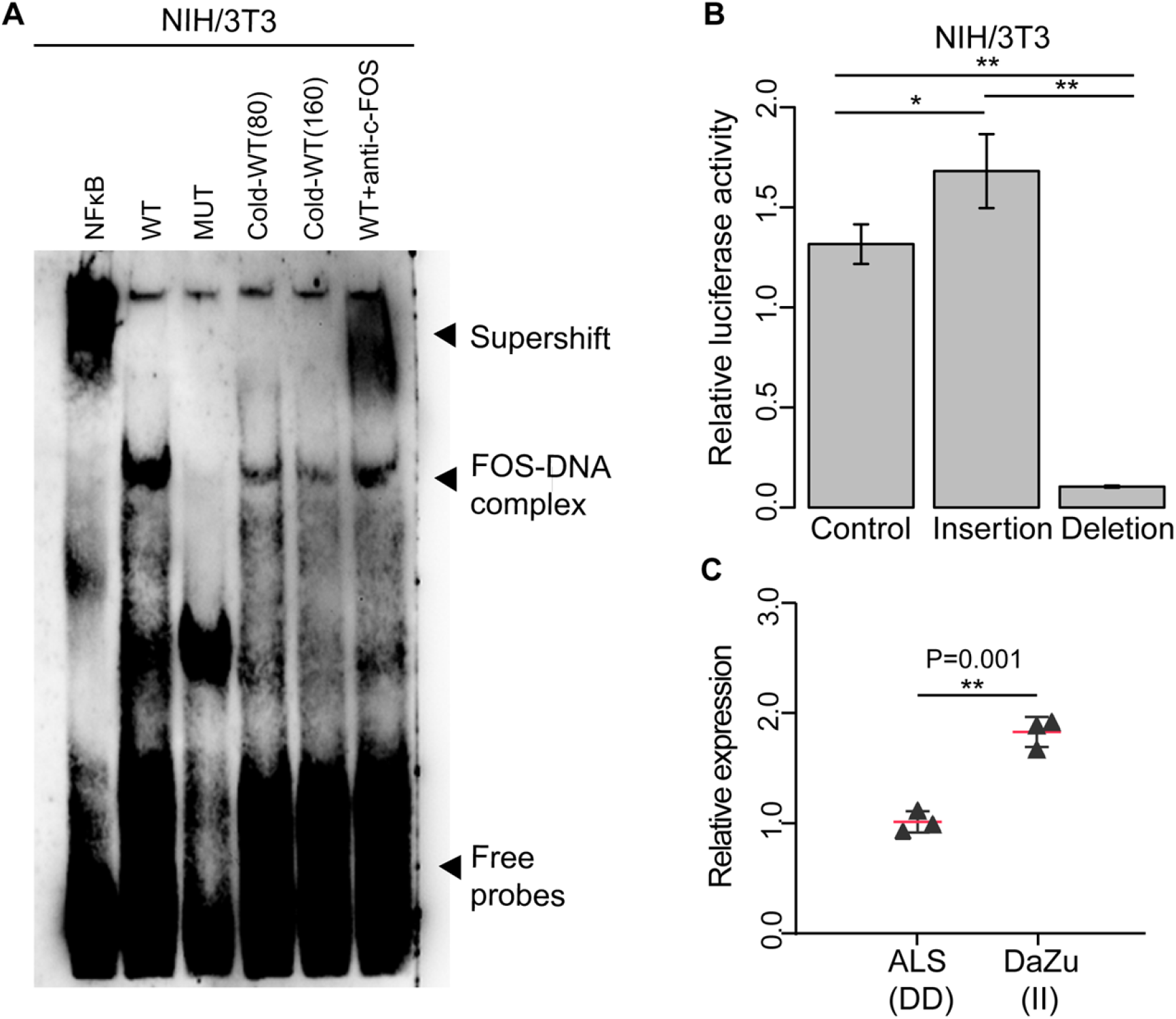
Validation of the indel variant in the *FGF5* gene. (A) Electrophoretic mobility shift assays (EMSAs) using the nuclear protein from NIH/3T3 cells. NFκB acts as a positive control (lane 1). WT and MUT represent the probe containing the wildtype FOS binding site and the probe excluding the FOS binding site (lane 2 and 3), respectively. The cold competitions of the protein complex formation by 80 and 160 fold over that of wildtype probe (lane 4 and 5). The clear supershift with anti-c-FOS antibody mixtured to wildtype react (lane 6). (B) Dual-luciferase activity assay of the NIH/3T3 cell lysates cotransfected with the pGL4.74 internal reference plasmid and the pGL4.23 empty vector as control, the pGL4.23 recombinant plasmids of the insertion or the deletion variant. (C) The qPCR gene expression of the *FGF5* gene in the skin of ALS (Alashan cashmere goat) and DaZu black goats (non-cashmere goat). * and ** displayed the statistical significance of *P-values* <0.05 and 0.01, respectively.

To confirm our hypothesis, we constructed dual-luciferase recombinant plasmids either including (pGL4.23-ins) or excluding (pGL4.23-del) the indel fragment. These two recombinant vector plasmids and an empty vector were each transfected into NIH/3T3 cells together with an internal luciferase control (pGL4.74) to measure the luciferase activities. We observed significantly higher luciferase activity in cells expressing the indel fragment than both empty cells and cells expressing pGL4.23-del (**Fig 5B**). This assay suggested that the indel fragment of the *FGF5* gene functions as an enhancer to which certain transcription factors specifically bind to upregulate *FGF5* gene expression.

We next examined whether the indel mutation could alter the transcriptional response to cashmere length using RT-qPCR assays. The mRNA expression level of cashmere goats carrying the deletion was significantly decreased compared to that of non-cashmere goats carrying the insertion (*P*<0.01, **Fig 5C**). Thus, the results of the gel shift experiment, dual-luciferase assay and RT-qPCR assays confirmed that the deletion variant disrupts the binding of transcription factors (e.g. FOS) and leads to lower expression of the *FGF5* gene in the skin of cashmere goats, while the insertion variant serves as an enhancer element that amplifies the transcriptional activity mediated by *FGF5* in non-cashmere goats.

## Discussion

In this study, we sequenced the genomes of 115 goats from 15 breeds originating from China, Nepal and Pakistan. The genome data set allowed us to identify a total of ~17.5 million SNP variants, which helped us reveal the genetic diversity and population structure of these goats. In this study, most of goat breeds showed a higher diversity than others, such as the Tibetan cashmere goats, dairy goats, Erlangshan cashmere goat and Xiangdong black goat. However, the Liaoning, Alashan and Arbus cashmere goats showed lower diversity than other Chinese goats, in line with previous work [10]. The three cashmere goat breeds are famous worldwide for their fine, long fibres. This fact indicated that these breeds may have been subject to stronger intensive selection for cashmere growth. An interspecies comparison showed that Pakistani goats had a lower diversity than others. Notably, similar to previous work based on the goat 50K SNP chip [10], the Bugi Toori (BTR) goat breed showed the lowest genetic diversity and a great differentiation from other Pakistani goat breeds. There may have been a historical bottleneck in history or an in-flight phenomenon in the BTR goat breed (**S1 and S2 Figs, S3 Table**).

Population structure analysis revealed that all the goats evaluated in this study were divided into six clusters, namely, dairy goat, southern Chinese goat, northern Chinese goat, Tibetan goat, Nepalese goat and Pakistani goat. When potential migration edges were added to the ML-TreeMix tree, gene exchange between the wild goats and Nepalese goats as well as Pakistani goats was detected among the clusters. This result may indicate that the local goats had a hybridization event with wild goats in the past. However, we observed migration edges between the Xiangdong black goat and Liaoning cashmere goat, as well as the dairy goat and Nepalese goat. There is a lower possibility of genetic admixture or introgresssion between the breeds because of the geographically distance between their inhabited regions. Therefore, to determine whether the southern Chinese goats underwent gene exchange with northern Chinese goat or Nepalese goats, the inclusion of more southern Chinese goat breeds is required. (**Fig 1B-1D, S3-S5 Figs**).

Compared with other domestic animals, goats are more adaptable to extreme environments. In China, cashmere goats are mainly distributed in the northern China and the Tibetan Plateau, where they have adapted well to the cold environment. More importantly, the fluff produced by cashmere goats provides good, warm materials for the native population. Therefore, different cashmere traits are continuously formed under natural and artificial selection. This study compared the genomes of cashmere goats including those from habitats in Liaoning, Inner Mongolia, Tibetan areas and Nepalese highland areas bordering the Tibetan region with various non-cashmere goats from different areas. Scanning the genome of cashmere goat breeds for signatures of positive selection revealed the *FGF5* gene among the top candidates (**Fig 2**). The *FGF5* gene participates in the FGF pathway, which plays a central role in hair growth. Studies on the *FGF5* gene demonstrated the relation to coat hair length in mice [29], dogs [29], cats [31], humans [32], donkeys [33] and alpacas [34]. The same selection target has been described in a number of cashmere goat investigations [10,35]. In addition, disruption of the *FGF5* gene via the CRISPR/Cas9 system in cashmere goats increased the number of secondary hair follicles and enhanced the fiber length [36]. Previous studies have found a few SNP variants of the *FGF5* gene that may be associated with hair length, including a missense SNP (c.284G> T) in dogs, four SNPs (c.194C>A, c.182T>A, c.474delT and c.475A>C) in cats, two SNPs (c.433_434delAT and c.245G> A) in donkeys and a missense SNP (c.499C>T) in alpacas [30,31,33,34]. Recently, one SNP (c.253G>A) in the 5’-UTR of *FGF5* resulted in a start codon that could lead to a premature/dysfunctional protein in Tibetan cashmere goats [35].

At the molecular level, our work did not reveal any missense SNPs in the exons of *FGF5* but instead revealed a significant indel variant in the region downstream of the *FGF5* gene locus (**Fig 3**). Interestingly, the result of the expanded population verification showed that the 505-bp indel variant was significantly separated in cashmere goats versus non-cashmere goats. The cashmere goats mainly exhibited a deletion mutation (> 0.9), whereas non-cashmere goats mainly exhibited an insertion mutation (~ 0.8). Some of the goat breeds have a small number of hybrids, such as the Nepalese lowland goat, Xiangdong black goat and Laoshan dairy goat, which may be caused by crossbreeding or altitude factors. Therefore, this result indicated that the indel variant can serve as a genetic marker for the cashmere growth trait. Furthermore, the indel mutation was found to contain a conserved binding site for the FOS transcription factor located in the mutation array and to be highly conserved in various mammals (**Fig 4**). In humans, the mutation is located downstream from the *FGF5* locus and has been identified as an enhancers according to the FANTOM5 Human Enhancers database (http://slidebase.binf.ku.dk/human_enhancers/). Therefore, it is speculated that the indel variant plays a potential enhancer role in *FGF5* gene transcription.

Thus, an electrophoretic mobility shift assay (EMSA), and a novel dual-luciferase reporter assay based on the expression of firefly and Renilla luciferase and mRNA expression levels of *FGF5* in goats were performed to explore the relevance of the indel variant to the *FGF5* gene. EMSA is a powerful tool for evaluating DNA-protein or RNA-protein interaction and is often used to detected the activated transcription factors (TF) that bind with DNA or RNA in the nucleus [37]. EMSAs based on NIH/3T3 nuclear extracts revealed that the protein complex bound to the biotin-probe containing the wildtype FOS binding site, but did not bind to the probe that contained the mutant FOS binding site. Efficient competition for protein complex formation was observed with the inclusion of a wildtype cold probe, and a clear supershift occurred when anti-c-FOS antibody was added, which further confirmed that the FOS transcription factor can specifically bind to the indel variant. The c-FOS protein is a member of the FOS protein family [38]. The dual-luciferase reporter gene assay is widely used to study promoter activity, transcription factors, intracellular signaling, protein interactions [39], miRNA regulation [40], and target site recognition [41]. The dual reporter gene assay based on firefly (*Photinus pyralis*) and sea kidney (*Renilla reniformis*, also known as marine pansy) luciferases can improve experimental accuracy by normalizing results and reducing technical differences [42]. The results of this assay showed that the insertion mutation can significantly enhance promoter transcription and increase gene expression thereby verifying its enhancer function. Finally, we detected significant differences in the expression levels of the *FGF5* gene from skin tissues of cashmere goats compared with non-cashmere goat, further confirming that the insertion variant may serve as an enhancer by binding to a transcription factor to result in increased transcription of its upstream *FGF5* gene target (**Fig 5**).

In conclusion, our study provides a whole-genome sequence analysis of Chinese, Nepalese and Pakistani goat breeds. It includes a total of 115 individual genomes spread across 15 goat breeds. The phylogenetic relationship of the 115 individuals revealed genetically distinct clusters according to their geographic locations, but genetic admixture or introgression may have occurred between Chinese and Nepalese goats. Genomic regions showing signatures of positive selection in cashmere goats revealed that the *FGF5* gene was the top candidate for the cashmere growth trait. Genotyping data from a large panel of 288 cashmere and non-cashmere goats revealed that a 505-bp indel variant, located downstream from the *FGF5* gene, is strongly associated with the cashmere length phenotype; furthermore, the deletion fragment reached close-to-fixation (~90%) frequencies in cashmere goats. Functional assays demonstrated that the insertion variant may act as an enhancer by binding to transcription factor, ultimately causing increased transcription of the upstream *FGF5* gene target. Our study provides useful information for the sustainable utilization and improved conservation of goat genetic resources. The valuable genetic marker that we identified will contribute to cashmere goat breeding to improve cashmere growth in the future.

## Materials and Methods

All sample collection were approved by the Animal Welfare and ethics Committee of Institute of Animal Science, Chinese Academy of Agricultural Sciences (Permit number: IAS2019-61).

### Sample information

In this study, We collected a total of 91 goats representing 44 Chinese native goats, 16 Nepalese goats and 31 Pakistani goats for whole-genome sequencing. In addition, we downloaded genomic dataset of 24 Chinese goats from the Sequencing Read Archieve (https://www.ncbi.nlm.nih.gov/) under accession code PRJNA338022. 51 of the total 115 goats are cashmere goats, including 8 Liaoning (LiNi), 6 Arbus (ABS), 7 Erlangshan (ELS), 3 Alashan (ALS), 10 Tibetan Bange (TBG), 10 Tibetan Ritu (TRT) in China and 7 Nepalese highland (NPH) in Nepal. 64 native goats produce little cashmere, including 10 Toggenburg dairy (TGB), 7 Laoshan dairy (LaSh), 7 Xiangdong black (XiDo) in China, 9 Nepalese lowland (NPL) in Nepal, 6 Bugi Toori (BTR), 9 Kamori (KMR), 11 Pateri (PTR) and 5 Tapri (TPR) in Pakistan (**S1 Table, Fig 1A**). A minimum of two separate flocks were sampled for each breed or location, and parent/offspring pairs were excluded.

### Whole genome sequencing analysis

DNA extraction was conducted by Wizard^®^ Genomic DNA Purification Kit (Promega). About 3μg of genomic DNA from each collected sample was sequenced on Illumina HiSeq 2000 instruments at BerryGenomics Company (Beijing, China). The 350bp sequencing library with paired-end sequencing was constructed using Illumina’s standard protocol. At least 5x genome coverage and no less than 15Gb sequencing data were gained per individual. After trimming low-quality bases and adapter sequences, the clean reads were aligned against the latest goat reference genome assembly ARS1(GCF_001704415.1) [43] using mem algorithm in Burrows-Wheeler Aligner (BWA) software [44,45]. Then the mapping results were converted to BAM format by SAMtools (Version: 1.1) [46]and sorted by SortSam tools in Picard packages (picard.sourceforge.net, Version: 1.86). Only properly paired reads both aligned to the reference were retained for subsequent analysis (**S2 Table**). The BamCoverage (https://github.com/BGI-shenzhen/BamCoverage/) was used to compute the coverage and depth of sequence alignments, with the “statistics Coverage” parameter.

### Variant calling

The program Genome Analysis Toolkit (GATK) [47]and SAMtools v0.1.19 [48] was used to identify SNPs, short insertions and deletions (indels). Reads were realigned around indels using the Realigner Target Creator and Indel Realigner tools from GATK, before calling SNPs with the GATK Unified Genotype and SAMtools mpileup modules, separately. SNPs were retained if matching the following five criteria: (1) the SNP confidence score (QD) was greater than or equal to 20; (2) the Phred-scaled P-value of the Fisher’s exact test to detect strand bias (FS) was inferior to equal to 10; (3) the Z-score of the Wilcoxon rank sum test of Alt vs. Ref read position bias (ReadPosRankSum) was greater or equal to −8; (4) the Qualscore of each individual SNP was larger-than-average; (5) SNPs showed only two possible alleles and a minimal allele frequency of 5%. The variants with sequence coverage and base-level values lower than the average of all sites were filtered. The 115 individual SNP VCF files were combined into the merged dataset of 17,534,538 autosomal SNPs and this merged SNP dataset was further phased to impute its own missing positions using BEAGLE software [49,50].

### Annotation

SNP variants were classified into protein coding regions (overlapping a coding exon), 5’UTRs and 3’UTRs (overlapping untranslated region), intronic regions (overlapping with an intron), or intergenic regions using the goat genome GTF file downloaded from Ensembl 94 (ftp://ftp.ensembl.org/pub/release-100/gtf/capra_hircus/) and the SNPEff software (Version: 4.0) [51]. SNPs located within protein coding regions were further binned into synonymous and non-synonymous SNPs **(S3 Table**).

### Diversity analysis

The within-population genetic diversity for goat populations was assessed using the filtered SNPs and various metrics, including observed (Ho) and expected heterozygosity (He) **(S4 Table)**. The runs of homozygosity (ROH) for each goat population, including the number of ROHs and the total size within ROHs for each individual, were calculated by the command of “--homozyg-window-snp 50 --homozyg-window-het 1 --homozyg-kb 500 −homozyg-density 1000” using the program PLINK v1.90b [52]. Linkage disequilibrium (LD) was computed for each population via the squared correlation coefficient (*r^2^*) between pairwise SNPs by the command of “-MaxDist 500 -MAF 0.005 -Het 0.9 −Miss 0.25” using the software PopLDdecay (https://github.com/BGI-shenzhen/PopLDdecay)

### Phylogenetic analysis

All SNPs were pruned using PLINK (Version:1.90b) and considering window sizes of 1000 variants, a step size of 5, and a pairwise r^2^ threshold of 0.5 (--indep-pairwise 1000 5 0.5). The principal component analysis (PCA) was carried out using the GCTA 1.91 software [53]. The neighbor-joining tree [13] was constructed using PHYLIP 3.68 (evolution.genetics.washington.edu/phylip.html). MEGA7 software [54] was used to visualize the phylogenetic trees. The population structure was examined via calculating Cross Value with an expectation maximization algorithm implemented in the software ADMIXTURE [55]. The number of assumed genetic clusters K ranged from 2 to 7. The population-level admixture analysis was conducted by TreeMix v.1.12 [12]. The program inferred the ML tree for 15 goat breeds (117 individuals) and an outgroup (wild goat). The command was ‘-I input bootstrap -k 10000 –root outgroup -o output’. From one to 6 migration events were gradually added to the ML tree, and the command was ‘-i input -bootstrap -k 10000 -m migration events -o output’.

### Selective sweeps

We scanned the cashmere goat genome for signatures of positive selection by combing three selection signature tests of the population-differentiation statistic (*F*_ST_) [12], the relative nucleotide diversity (θ_π_ ratio, θ_π-Noncash_/θ_π-Cash_) [57] and the transformed heterozygosity score (ZHp). Genomic evidence for positive selection in response to cashmere growth was evaluated by contrasting differentiation indices between the cashmere goats versus the other goats. *F*_ST_ and nucleotide diversity (θ_π_) were calculated by VCFTools [58]. The window-based ZHp approach was calculated as previously described [59]. Each test was based on a 100-kb window with 10-kb increment. We considered top 1% level for empirical percentile (*F*_ST_>0.153, θ_π_ ratio>1.547, |ZHp| >3.210) windows as candidate outliers in strong selective sweeps. To annotation candidate genes harbored in these selective regions, we used Rscript to map genes in selective windows. The overlapping windows shared by top 5% highest all three tests were considered as conservative candidate selection targets and were further annotated by the genomic database BioMart (http://www.biomart.org/). To detect the genomic loci that are associated with cashmere length around the *FGF5* gene, we also calculated the θ_π_ ratios, Tajima’*D* [60] and pairwise *F*_ST_ in 2 Mb windows between the cashmere breeds and the non-cashmere breeds. The Gene Ontology (GO) enrichment analysis of the annotated candidates were performed by using both the online G:profiler.

### Validation in the extended population

In order to predict functional candidates, this 505 bp indel variant returning the most significant signature was classified according to their evolutionary conservation scores among other mammals. Primers were designed according to the indel region of FGF5 gene: FGF5-indel-F: 5’-GGTGATAAGCCACACGTTCAAA-3’, FGF5-indel-R: 5’-TGGCTGTGATCAAACTTACAACC-3’. The indel region was genotyped by PCR amplification using the reaction condition of the 5-min pre-denaturation, 30s-denaturation, 30s-annealing and 45s-extension for 40 cycles. The genotype results were visualized by agarose gel electrophoresis. The indel of *FGF5* gene were successfully genotyped in the extended population of 288 goats, including 153 cashmere goats and 135 non-cashmere goats.

### Electrophoretic mobility shift assay

The crude nuclear protein was extracted from NIH/3T3 cells using the Nuclear/Cytoplasmic Protein Extraction Kit (SINP001, Viagene) and protein concentration was determined by the Enhanced BCA Protein Assay Kit (CHEM001, Viagene). The oligonucleotide probe of the wild allele was 5’-ATGACTC**TGAGTCA**GTCTCCTCC-3’, while the oligonucleotide probe of the mutant allele was 5’-ATGACTCGTCTCCTCC-3’. The probes were synthesized by Viagene Biotech company. EMSA was performed using a non-radioactive EMSA kit (SIDET101, Viagene) with biotin-probes, according to the user’s manual instruction. Briefly, 4 μg nuclear protein was incubated with poly dI:dC for 20 min at room temperature in binding reaction buffer. Then biotin-probe was added to and incubated with the mixture at room temperature for at least 20 min. The reaction mixtures were separated by electrophoresis by 8% non-denaturing polyacrylamide gel in 0.5× Tris-borate-EDTA buffer at 120V for 1h. The gel was transferred onto a pre-soaked nylon-membrane at 390 mA for 40min and afterward, the energy of 800 mJ is applied for crosslinking DNA to nylon membrane using CL-1000 Ultraviolet Crosslinker (UVP, UK). Finally, the complexes bands were visualized by chemiluminescent detection. Competition reaction with a 80-fold and 160-fold molar excess of unlabeled oligonucleotide were performed to confirm the specificity of the DNA-protein complex. For the supershift experiment, 2μg of anti-c-fos antibodies (sc-8047X, Santa Cruz) were added to the mixture.

### Dual-luciferase reporter assay

NIH/3T3 cells were propagated in the medium of Roswell Park Memorial Institute 1640 (RPMI 1640), supplemented with 10% heat-inactivated fetal bovine serum and penicillin (0.2 U/ml)/streptomycin (0.2 μg/ml)/L-glutamine (0.2 μg/ml) (Gibco, USA). The 772-bp and 267-bp fragment of *FGF5* indel region were cloned into the pGL4.23 vector (Promege, USA) expressing Firefly luciferase gene, respectively. We thus generated two recombinant vectors pGL4.23-ins (containing 505-bp insertion) and pGL4.23-del (containing the deletion). Each plasmid was co-transfected into NIH/3T3 cells with the internal control vector pGL4.74 expressing Renilla luciferase gene by Lipofectamine™ 3000 (Invitrogen, America) according to the manufacturer’s instruction. The firefly luminescence signal (FiLuc) and Renilla luciferase signal (hRLuc) of NIH/3T3 cells were measured for each transfection on a multi-function microplate reader (Tecan Infinite 200 Pro) using the Dual-Luciferase Reporter Assay System (E1910, Promega) after 24h transfection.

### RT-qPCR quantification

The total RNA was extracted from Inner Mongolia Alashan cashmere goats and Dazu black goats using RNA extraction Kit (Promega), and RNA quality and concentration were measured on an Agilent 2100 Bioanalyzer (Germany). The RIN value of samples greater than 8.0 were used for RT-qPCR analysis. The cDNA was synthesized using PrimeScript RT reagent kit with gDNA Eraser (Takara, Dalian, China) in a 20 μl reaction mixture following the manufacturer’s instruction. The expression levels of FGF5 gene was normalized against UBC reference genes [61,62]. The primers used in the RT-qPCR experiment were showed in **S11 Table**. The RT-qPCR was performed using TB Green Premix Ex Taq (Takara, Dalian, China). The qPCR reaction program was set as follow: 95 °C 30 s, 40 cycles of 5 s at 95 °C and 34 s at 60 °C. The qPCRs were run of both technical and biological replicates (n=3) using an ABI7500 sequence detection system (Applied Biosystems by Life Technologies, Darmstadt, Germany). Fold expression changes were determined using a standard 2^−ΔΔCT^ method that compares C_T_ (cycle threshold) values of a reference gene to the gene of interest for the ΔC_T_ calculation and compares the ΔC_T_ value of a reference sample with the sample of interest for the ΔΔC_T_ calculation [63].

## Acknowledgments

The authors are grateful to all goat owners and breeding organizations who donated samples. We thank members of the Nextgen project for sharing their data. We thank the National Germplasm Center of Domestic Animal Resources of IAS.

## Supporting information

**S1 Fig. Distribution of mean size and number of ROH.**

TGB, Toggenburg dairy goat from Heilongjiang; LaSh, Laoshan dairy goat from Shandong; XiDo, Xiangdong black goat from Hunan; LiNi, Liaoning cashmere goat from Liaoning; ABS, Arbus cashmere goat; ELS, Erlangshan cashmere goat; ALS, Alashan cashmere goat from Inner Mongolia; TBG, Tibetan Bangor cashmere goat; TRT, Tibetan Ritu cashmere goat from Tibet; NPH, Nepalese Highland goat; NPL, Nepalese Lowand goat; TPR, Tapri goat; KMR, Kamori goat; PTR, Pateri goat; BTR, Bugi Toori goat.

**S2 Fig. Decay of LD in the goat genome for each breed.**

**S3 Fig. PCA result of the first and third components of Chinese goats.**

**S4 Fig. Genetic population structure of the 115 goats conducted by Admixture.**

The length of each colored segment represents the proportion of the individual genome inferred from ancestral populations (K=2-7).

**S5 Fig. Migration analysis of 115 goats by Treemix software.**

(A-F) panels represent models of population affinities assuming 0-3 and 5-6 migration edges in TreeMix, respectively. The inferred migration weight is provided by the color of the arrow displayed.

**S6 Fig. The overlapped regions for selection signatures.**

The overlapped regions by top 5% highest *F*_ST_ and θ_π_ ratio (noncash/cash) and ZHp for cashmere goats.

**S1 Table. Population distribution.**

**S2 Table. Mapping coverage and depth.**

**S3 Table. SNP summary statistics in the goat breeds.**

**S4 Table. Genome-wide heterzygosity of goat breeds.**

**S5 Table. *F*_ST_ selection signatures with the windows by top 1% highest.**

**S6 Table. θπ selection signatures with the windows by top 1% highest.**

**S7 Table. ZHp selection signatures with the windows by top 1% highest.**

**S8 Table. Overlapped regions by top 5% highest *F*_ST_, θπ and ZHp selection signatures.**

**S9 Table. Enrichment analyses of GO terms.**

**S10 Table. Populations for validation.**

**S11 Table. Primers for qPCR.**

**S12 Table. θπ values of genomic regions around gene *FGF5*.**

**S13 Table. Tajima’*D* values of genomic regions around gene *FGF5*.**

**S14 Table. *F*_ST_ values of genomic regions around gene *FGF5*.**

**S15 Table. Depth values of genomic regions around gene *FGF5*.**

